# Deficiencies of Runx3 and tissue-resident CD4^+^ intestinal epithelial lymphocytes link intestinal dysbiosis and inflammation in mouse and human spondyloarthropathy

**DOI:** 10.1101/2022.07.28.501834

**Authors:** Zaied Ahmed Bhuyan, Muralidhara Rao Maradana, M. Arifur Rahman, Ahmed Mehdi, Anne-Sophie Bergot, Davide Simone, Aimee L Hanson, Hendrik Nel, Jose Garrido-Mesa, Marya El-Kurdi, Tony Kenna, Paul Leo, Linda Rehaume, Matthew A Brown, Francesco Ciccia, Ranjeny Thomas

## Abstract

**Objective:** Disturbances in immune regulation, intestinal microbial dysbiosis and intestinal inflammation characterize ankylosing spondylitis (AS), which is associated with *RUNX3* loss-of-function variants. ZAP70^W163C^ mutant (SKG) mice have reduced ZAP70 signaling, spondyloarthritis and ileitis. At intestinal epithelial interfaces, lamina propria Foxp3^+^ regulatory T cells (Treg) and intraepithelial CD4^+^CD8αα^+^TCRαβ^+^ lymphocytes (CD4-IEL) control inflammation. TGF-β and retinoic acid (RA)-producing dendritic cells are required for induction of Treg and for CD4-IEL differentiation from CD4+ conventional or Treg precursors, with upregulation of Runx3 and suppression of ThPOK. We investigated Treg, CD4-IEL, ZAP70 and Runx3 in SKG mice and AS patients.

**Methods:** We compared ileal Treg and CD4-IEL numbers and differentiation in BALB/c and SKG mice, and with ZAP70 inhibition, and related differentially-expressed genes in terminal ileum to ChIP-seq-identified Runx3-regulated genes. We compared proportions of CD4-IEL in ileum and CD4^+^8^+^ T cells in blood of AS patients and healthy controls.

**Results:** ZAP70^W163C^ or ZAP70 inhibition prevented intestinal CD4-IEL but not Foxp3^+^ Treg differentiation in context of TGF-β and RA in vitro and in vivo, resulting in Runx3 and ThPOK dysregulation. CD4-IEL frequency and expression of tissue resident memory T-cell and Runx3-regulated genes was reduced in SKG intestine. Multiple under-expressed genes were shared with risk SNPs identified in human spondyloarthropathies. CD4-IEL were decreased in AS intestine. Double-positive T cells were reduced and Treg increased in AS peripheral blood.

**Conclusion:** High-affinity TCR-ZAP70 signalling is required for Runx3-mediated intestinal CD4-IEL differentiation from Treg. Genetically-encoded relative immunodeficiency of T cells underpins poor intestinal barrier control in mouse and human spondyloarthropathy.

**What is already known about this subject?:** Ankylosing spondylitis (AS) is associated with RUNX3 loss-of-function variants.

Capacity of the AS T cell receptor repertoire to expand in response to infectious antigens is reduced.

Foxp3^+^ regulatory T cells (Treg) are increased in AS intestine.

ZAP70W163C mutant (SKG) mice have reduced ZAP70 signaling, spondyloarthritis (SpA) and ileitis.

Intestinal epithelial Foxp3^+^ Treg and CD4^+^CD8^+^ cytotoxic lymphocytes (CD4-IEL) control local inflammation. CD4-IEL differentiate from Treg, with upregulation of Runx3 and suppression of ThPOK transcription factors.

**What does this study add?:** High-affinity TCR-ZAP70 signalling is required for Runx3-mediated intestinal CD4-IEL differentiation from Treg

Intestinal CD4-IEL and circulating CD4^+^CD8^+^ T cells are reduced in AS while circulating Treg are increased. Impaired CD8 expression may be correctible by TNF inhibition in AS CD4^+^ T cells.

Deficiencies of Runx3 and tissue-resident CD4-IEL link intestinal dysbiosis and inflammation in mouse and human SpA.

**How might this influence clinical practice or future developments?:** Genetically-encoded relative T immunodeficiency underpins poor intestinal barrier control in SpA

## Introduction

Spondyloarthropathies (SpA) comprise a group of common, strongly heritable inflammatory diseases including ankylosing spondylitis (AS), psoriatic arthritis and arthritis associated with inflammatory bowel disease (IBD). Shared polymorphisms associated with SpA are in genes related to adaptive and innate immune function, infection control, antigen processing and presentation, T cell signaling and differentiation, IL-23 signaling, autophagy and NF-κB pathways [1]. Loss of function *RUNX3* variants are associated with AS and psoriatic arthritis risk [1, 2]. In AS, disturbances in immune regulation are associated with intestinal microbial dysbiosis and subclinical intestinal inflammation, indicative of impaired intestinal immune homeostasis [3-6]. In the 46% of AS patients with intestinal inflammation, arthritis activity correlated with flares in bowel disease [7]. Furthermore, *RUNX3* risk SNPs have been associated with higher AS disease severity [8].

The normal gut harbours CD4^+^ T cells with effector and regulatory function. Foxp3^+^ regulatory (Treg) cells limit inflammatory activation of effector and autoreactive T cells to curtail autoimmune tissue inflammation. Treg also suppress innate immune cells, limiting innate bacterial activation at epithelial interfaces. Treg are either agonistically selected in the thymus (tTreg) or induced from CD4^+^ T cells in peripheral organs (pTreg). pTreg are particularly likely to develop in the small intestine and draining mesenteric lymph nodes in response to antigen presented by specialized CD103^+^CD11b^-^ dendritic cells (DCs) [9-11]. Small intestine also contains TCRαβ^+^CD4^+^CD8^+^ cytotoxic intra-epithelial T cells (CD4-IEL) with anti-inflammatory and regulatory function. Mature CD4^+^ T cells and Treg are converted to CD4-IEL in the intestinal epithelium [12-14]. This conversion is promoted in MHC class I-restricted T cell-associated molecule (CRTAM)-expressing precursor CD4^+^ T cells [15] again through recognition of antigen presented by specialized CD103^+^CD11b^-^ intestinal DCs [16]. Signals from TGF-β and retinoic acid (RA) critically determine the fate of CD4^+^ T cell differentiation in the periphery by transcriptionally reprogramming precursor CD4^+^ T cells to differentiate into pTreg and CD4-IEL [11, 17, 18]. The regulatory and cytotoxic function of CD4-IEL may compensate for Treg deficiency in the gut, providing an additional level of intestinal inflammatory control [13, 19]. In this TGFβ/RA context, Thpok, the CD4^+^ T-cell fate-determining zinc-finger repressor decreases and CD8^+^ T cell fate-determining Runx3 increases in CD4^+^ T cells, promoting cytolytic and innate-like functions, CD8αα expression and CD4IEL cell development [12]. Components of the nucleosome remodeling and deacetylase (NuRD) complex interact with Thpok in CD4^+^ T cells [20], and this recruitment of NuRD complex is central to repression of Runx3 gene expression. In CD8^+^ T cells, an AS-risk allele of RUNX3 was shown to bind components of the NuRD complex and Aiolos more strongly than the protective allele [21].

Antigen recognition is required for TGF-β and RA-induced modulation of CD4^+^ T cells to CD4-IEL. Intestinal microbiota are essential for CD4-IEL cell development, as CD4-IEL induction was abrogated in antibiotic-treated mice and germ free mice [13, 19]. Although the antigen specificity of CD4-IEL in conventionally-raised mice is not restricted, clonal CD4-IEL developed from CD4^+^ T cells, possibly driven by *Lactobacillus reuteri*, in an AhR-dependent manner [22]. Few CD4-IEL were present in naïve ovalbumin (OVA)-specific T cell receptor (TCR) transgenic mice. However, they were induced after OVA feeding [12]. Thus intestinal antigens derived from the microbiota and the diet support CD4-IEL development in the intestine.

CD4^+^ T cells recognize antigen with varying degrees of affinity: high affinity TCR activation promotes Treg cell development in the thymus and pTreg development is increased by presentation of high affinity T cell epitopes or decreased intraclonal competition [19, 23]. However, it is not clear whether TCR affinity influences pTreg and CD4-IEL cell development from precursor CD4^+^ T cells in the intestine. BALB/c ZAP-70^W163C^ mutant (SKG) mice have reduced TCR affinity, which skews thymic selection towards auto-reactive CD4^+^ T cells with a propensity to become Th17 cells [23]. SKG mice raised in specific pathogen free (SPF) conditions injected with microbial 1,3-glucan (curdlan) develop interleukin (IL)-23/IL-22/IL-17 axis-dependent SpA, including arthritis of spine and joints, and Crohn’s-like ileitis associated with dysbiosis [24]. Here we show differential capacity of ZAP70 to regulate CD4^+^ T cell response to differentiation signals for Foxp3^+^ pTreg and CD4-IEL in the intestine of mice and humans with SpA.

## Materials and methods

### Patients

Two HLA-B27+ AS patients, who fulfilled modified New York criteria and had active disease (AS disease activity score ≥2), underwent ileocolonscopy independent of gastrointestinal symptoms. Two age and sex-matched HC without evidence of intestinal disease were undergoing ileocolonoscopy for diagnostic purposes. Peripheral blood mononuclear cells (PBMC) were obtained from 128 AS patients and 34 HC (cohort 1); and 24 AS patients not treated with TNF inhibitors (TNFi), 28 TNFi-treated AS patients and 40 age and sex-matched HC (cohort 2), as described [25]. Additional methods for mice, cell purification, flow cytometry, statistical analysis, T cell differentiation in vitro and in vivo, are provided in the supplemental material.

## Results

### Intestinal tissue resident CD4-IEL but not Foxp3^+^ Treg are diminished in SKG mice

Naïve SKG mice have intestinal dysbiosis associated with reduced capacity for IL-22 production by ILC3 in the small intestine and thus represent an excellent model of IBD susceptibility [24, 26]. To further explore intestinal immune regulation in naive SKG mice, we analyzed subsets of CD4^+^ T cells with regulatory capacity in the small intestine (SI). The proportion of Foxp3^+^ Treg among CD4^+^ T cells was increased in naïve SKG mice relative to BALB/c mice (Figure 1A). However, the number of SI CD4^+^ T cells, and hence of CD4^+^Foxp3^+^ Treg, was significantly lower than in BALB/c mice (Figure 1B). The proportion of Helios^-^ pTregs was comparable in the SKG and BALBc SI (Figure S1). CD4^+^ T cells were significantly lower among intestinal epithelial lymphocytes (IEL) and lamina propria lymphocytes (LPL) of SKG mice compared to BALB/c mice (Figure 1C). Granzyme-B^+^ single CD4^+^ T cells were also reduced in SKG mice, suggesting a generalized deficiency in cytotoxicity (Figure 1D). The IEL tissue resident memory marker CD103 was similarly reduced in SKG CD4^+^ T cells (Figure 1E). Loss of Thpok and concomitant acquisition of Runx3 determines the cytolytic gene signature and IEL phenotype in CD4-IEL, and IFN-γ-dependent T-bet regulates Thpok and Runx3 mutual expression by directly binding to their transcription regulatory regions [12]. In the SI, Thpok was increased and Runx3 decreased in SKG relative to BALB/c CD4^+^ T cells (Figure 1F). Consistent with previous studies [27], CD4^+^ T cells from SKG mice expressed higher levels of T-bet (Figure 1F). Thus, enhanced T-bet is insufficient to induce CD4^+^ T helper cell trans-differentiation to CD-IEL. These results indicate that SI of SKG mice lacks CD4-IEL associated with deficient Runx3 expression, but that Foxp3^+^ pTreg proportion is increased.

**Figure 1.**
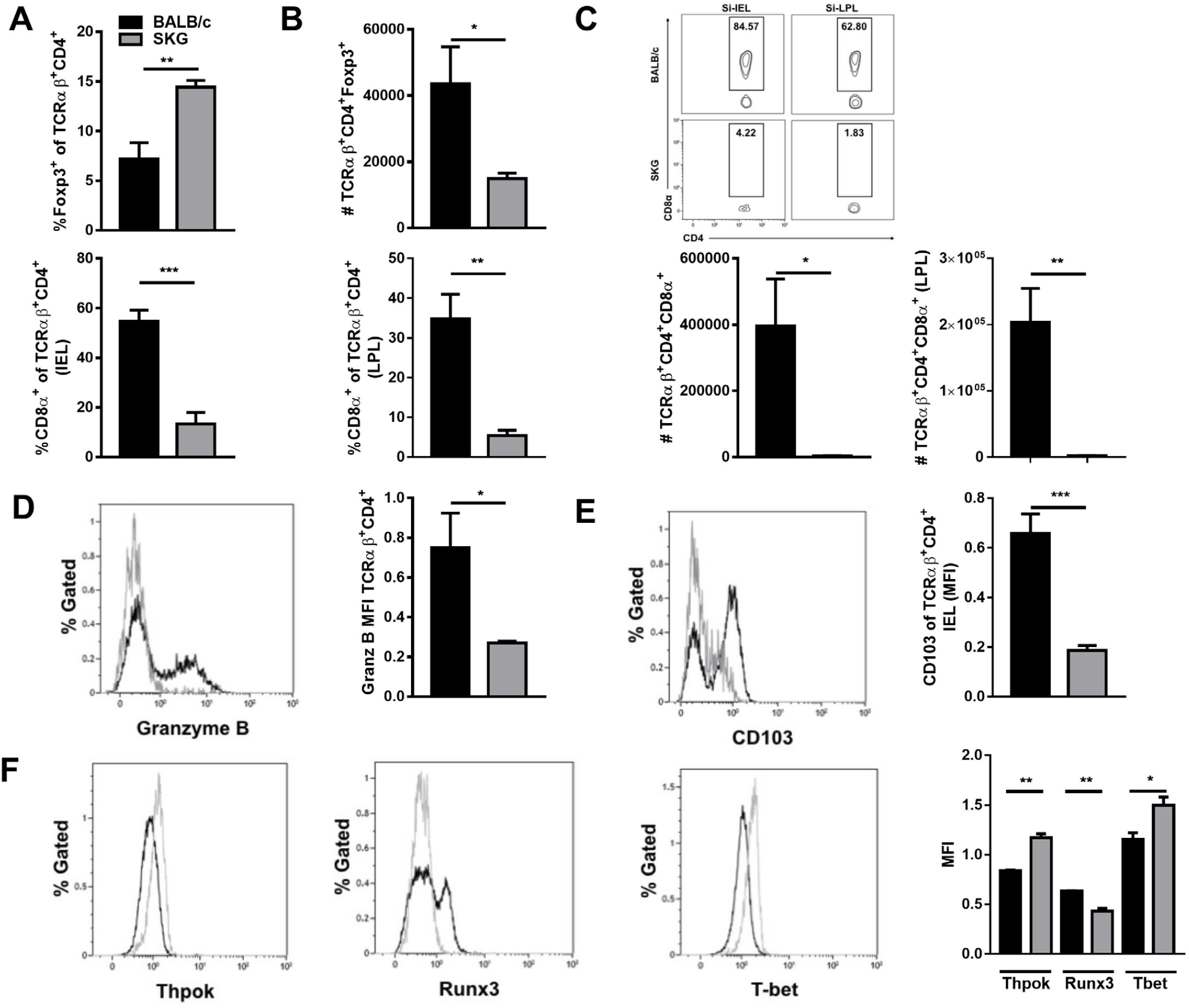
Intestinal tissue resident CD4+CD8+ T cells are diminished among intra-epithelial and lamina propria lymphocytes of SKG mice. Percentage Foxp3^+^ (**A**) total number of Foxp3^+^ (**B**) CD4^+^ T cells in the small intestine lamina propria of naïve BALB/c and SKG mice. **C**: Representative FACS plots and quantitation of CD4^+^CD8^+^T cells in small intestine epithelium (SI-IEL) and lamina propria (SI-LPL) of BALB/c and SKG mice. Percentage and number of CD4^+^CD8^+^ T cells in IEL (**D, F**) and LPL (**E, G**) respectively. Expression of intracellular granzyme B (**H, I**), surface CD103 (**J, K**) and intracellular transcription factors Thpok (**L, O**), Runx3 (**M, O**) and T-bet (**N, O**) by TCRβ^+^CD4^+^ T cells from intestinal epithelium from BALB/c and SKG mice. Data are represented as mean ± SEM. Representative of three separate experiments analyzing individual mice. n=5 per group. * *p*<0.05 ** *p*<0.01, t test.

### Differential expression of tissue resident memory T cell and Runx3-regulated genes in SKG ileum

In view of the markedly abnormal profile of CD4^+^ T cells in SI, we determined differentially-expressed genes in RNA extracted from terminal ileum of SKG and BALB/c terminal ileum to explore key dysregulated pathways. Tissue resident memory T cell-associated genes e.g. *Cd3e, Itgae, Ccl5*, and *Itgb7*, and antigen processing and presentation genes e.g. *H2-ab1, H2-Dma, H2-ea, Tap1*, and *H2-q8*, were among the top downregulated genes in SKG ileum (Table 1, Figure 2A). *Cx3cr1*, which is implicated in mucosal phagocyte differentiation and bacterial sampling and *Nkx2-3*, which is required for MadCAM-1-dependent lymphocyte homing to Peyer’s patches, were also downregulated in SKG ileum [16]. Over-expressed genes encoded intectin, a protein which accelerates intestinal epithelial cell apoptosis and IL-18, which promotes IBD and which is over-expressed in AS gut (Table 1)[28, 29]. Enriched GO terms include antigen processing and presentation, regulation of T cell activation and leukocyte cell adhesion (Supplementary Table 1). Since Runx3 expression is required for CD4-IEL development and was reduced in SKG CD4^+^ LPL (Figure 1G) we interrogated our DE genes for Runx3-regulated genes previously identified by ChIP-Seq [30]. Twenty five of the 68 (37%, odds ratio=2.13, Fisher exact p-value 0.004) of the DE genes are predicted to be Runx3-regulated (Supplementary Table 2, Figure 2B). Thus, consistent with the reduction in CD4-IEL and CD4^+^ T cell Runx3 expression, SKG SI is characterized by reduction of genes associated with antigen presentation to tissue resident memory T cells and regulation by Runx3. CD4-IEL cell development is mainly induced by cytokine signals, including TGF-β and IFN-γ [31]. In addition to these cytokines, RA produced by IRF8-dependent CD103^+^CD11b^-^ DCs influences CD4^+^ T cell trans-differentiation and maintenance of the IEL phenotype [12, 18]. Despite the overall reduction in antigen processing and presentation gene expression, SKG and BALB/c mice had comparable proportions and numbers of CD103^+^CD11b^-^ DCs (Figure 2C and Supplementary Figure 2) and MHC class-II expression by CD103^+^CD11b^-^ DCs was at higher levels in SKG than BALB/c mice (Supplementary Figure 2). Furthermore, receptors and signaling transcripts encoding these genes and genes involved in RA production such as *Aldh1a1* and *Aldh1a2* were similarly expressed in SKG and BALB/c ileum, indicating that neither the MHC class II signalling capacity of IRF8-dependent DCs nor RA/TGF-β cofactors were likely contributors to CD4-IEL cell deficiency in SKG mice.

**Table 1.** Differentially expressed genes in SKG relative to BALB/c ileum. Log fold change (FC), adjusted p-values (p_adj_) and gene description are shown.

| Gene | logFC | $p_{adj}$ | Gene description |
| --- | --- | --- | --- |
| H2-AB1 | -1.968 | 0.009026 | H-2 class II histocompatibility antigen, A beta chain |
| H2-AB1 | -1.896 | 0.009026 | H-2 class II histocompatibility antigen, A beta chain |
| ITGAE | -1.872 | 0.002153 | integrin, alpha E (antigen CD103, human mucosal lymphocyte antigen 1; alpha polypeptide) |
| ITGAE | -1.837 | 0.001049 | integrin, alpha E (antigen CD103, human mucosal lymphocyte antigen 1; alpha polypeptide) |
| CD177 | -1.836 | 0.001049 | CD177 molecule |
| H2-DMA | -1.491 | 0.03327 | Class II histocompatibility antigen, M alpha chain |
| CCL5 | -1.442 | 0.003549 | chemokine (C-C motif) ligand 5 |
| H2-EA | -1.381 | 0.03007 | H-2 class II histocompatibility antigen, E-U alpha chain |
| TAP1 | -1.289 | 0.003549 | transporter 1, ATP-binding cassette, sub-family B (MDR/TAP) |
| CTSW | -1.243 | 0.003549 | cathepsin W |
| CD59B | -1.224 | 0.01474 | CD59 molecule, complement regulatory protein |
| NUP210 | -1.196 | 0.004842 | nucleoporin 210kDa |
| MCM5 | -1.156 | 0.03239 | minichromosome maintenance complex component 5 |
| ITGB7 | -1.108 | 0.003194 | integrin, beta 7 |
| CCND2 | -1.071 | 0.04631 | cyclin D2 |
| NOXA1 | -0.9825 | 0.01645 | NADPH oxidase activator 1 |
| COTL1 | -0.94 | 0.009026 | coactosin-like 1 (Dictyostelium) |
| ACTA2 | -0.9231 | 0.01831 | actin, alpha 2, smooth muscle, aorta |
| CDT1 | -0.8805 | 0.03323 | chromatin licensing and DNA replication factor 1 |
| GVIN1 | -0.8702 | 0.04398 | GTPase, very large interferon inducible pseudogene 1 |
| TCIRG1 | -0.8664 | 0.01451 | T-cell, immune regulator 1, ATPase, H<sup>+</sup> transporting, lysosomal V0 subunit A3 |
| H2-Q8 | -0.8621 | 0.03716 | H-2 class I histocompatibility antigen, Q8 alpha chain |
| CD3E | -0.8288 | 0.01975 | CD3e molecule, epsilon (CD3-TCR complex) |
| MSC | -0.7953 | 0.009026 | musculin |
| ARHGAP30 | -0.7507 | 0.03716 | Rho GTPase activating protein 30 |
| GIMAP7 | -0.75 | 0.009026 | GTPase, IMAP family member 7 |
| PRMT5 | -0.6878 | 0.01187 | protein arginine methyltransferase 5 |
| NKX2-3 | -0.6845 | 0.04398 | NK2 homeobox 3 |
| NOXO1 | -0.6844 | 0.04398 | NADPH oxidase organizer 1 |
| SERPINA3G | -0.649 | 0.04944 | Serine protease inhibitor A3G |
| NSMCE1 | -0.6086 | 0.04398 | non-SMC element 1 homolog (S. cerevisiae) |
| CX3CR1 | -0.6028 | 0.0499 | chemokine (C-X3-C motif) receptor 1 |
| DGKG | -0.5958 | 0.04583 | diacylglycerol kinase, gamma 90kDa |
| DENND1C | -0.5955 | 0.02594 | DENN/MADD domain containing 1C |
| C1QTNF3 | -0.576 | 0.04583 | C1q and tumor necrosis factor related protein 3 |
| 1200009O22RIK | -0.5606 | 0.0469 | - |
| RAD54L | -0.5221 | 0.0499 | RAD54-like (S. cerevisiae) |
| ARRB2 | -0.513 | 0.03716 | arrestin, beta 2 |
| HIST1H3A | -0.4469 | 0.04398 | histone cluster 1, H3a |
| D6WSU176E | 0.5871 | 0.04398 | - |
| EPB4.1L3 | 0.6901 | 0.0499 | erythrocyte membrane protein band 4.1-like 3 |
| ITGB6 | 0.7005 | 0.03007 | integrin, beta 6 |
| FBXO32 | 0.7065 | 0.04807 | F-box protein 32 |
| TMEM128 | 0.8755 | 0.04398 | transmembrane protein 128 |
| TMBIM1 | 0.8937 | 0.04146 | transmembrane BAX inhibitor motif containing 1 |
| 4121402D02RIK | 0.9441 | 0.04398 | - |
| CHST8 | 0.9595 | 0.009026 | carbohydrate (N-acetylgalactosamine 4-0) sulfotransferase 8 |
| PTPRD | 0.9848 | 0.0469 | protein tyrosine phosphatase, receptor type, D |
| STK17B | 1.037 | 0.04398 | serine/threonine kinase 17b |
| DDR1 | 1.056 | 0.03716 | discoidin domain receptor tyrosine kinase 1 |
| LOC100046744 | 1.086 | 0.0365 | - |
| ARSA | 1.119 | 0.01187 | arylsulfatase A |
| ERRFI1 | 1.254 | 0.04398 | ERBB receptor feedback inhibitor 1 |
| LOC674706 | 1.257 | 0.007933 | - |
| MS4A10 | 1.313 | 0.004361 | membrane-spanning 4-domains, subfamily A, member 10 |
| PDLIM2 | 1.321 | 0.01054 | PDZ and LIM domain 2 (mystique) |
| SFRS5 | 1.381 | 0.04731 | serine/arginine-rich splicing factor 5 |
| HSD3B2 | 1.393 | 0.01637 | hydroxy-delta-5-steroid dehydrogenase, 3 beta- and steroid delta-isomerase 2 |
| NAIP1 | 1.448 | 0.03007 | Baculoviral IAP repeat-containing protein 1a |
| PIGT | 1.597 | 0.0499 | phosphatidylinositol glycan anchor biosynthesis, class T |
| 1300013J15RIK | 1.716 | 0.03716 | SLC47A1 |
| CYP2D22 | 1.816 | 0.002876 | cytochrome P450, family 2, subfamily D, polypeptide 6 |
| TRAPPC3 | 1.831 | 0.009026 | trafficking protein particle complex 3 |
| IL18 | 1.957 | 0.04398 | interleukin 18 (interferon-gamma-inducing factor) |
| SLC6A8 | 2.103 | 0.01645 | solute carrier family 6 (neurotransmitter transporter, creatine), member 8 |
| PPP3R1 | 2.104 | 0.0001553 | protein phosphatase 3, regulatory subunit B, alpha |
| ACOX2 | 3.009 | 0.0499 | acyl-CoA oxidase 2, branched chain |
| 2010109I03RIK | 11.22 | 9.506e-10 | Intectin. Ly6m |

**Figure 2.**
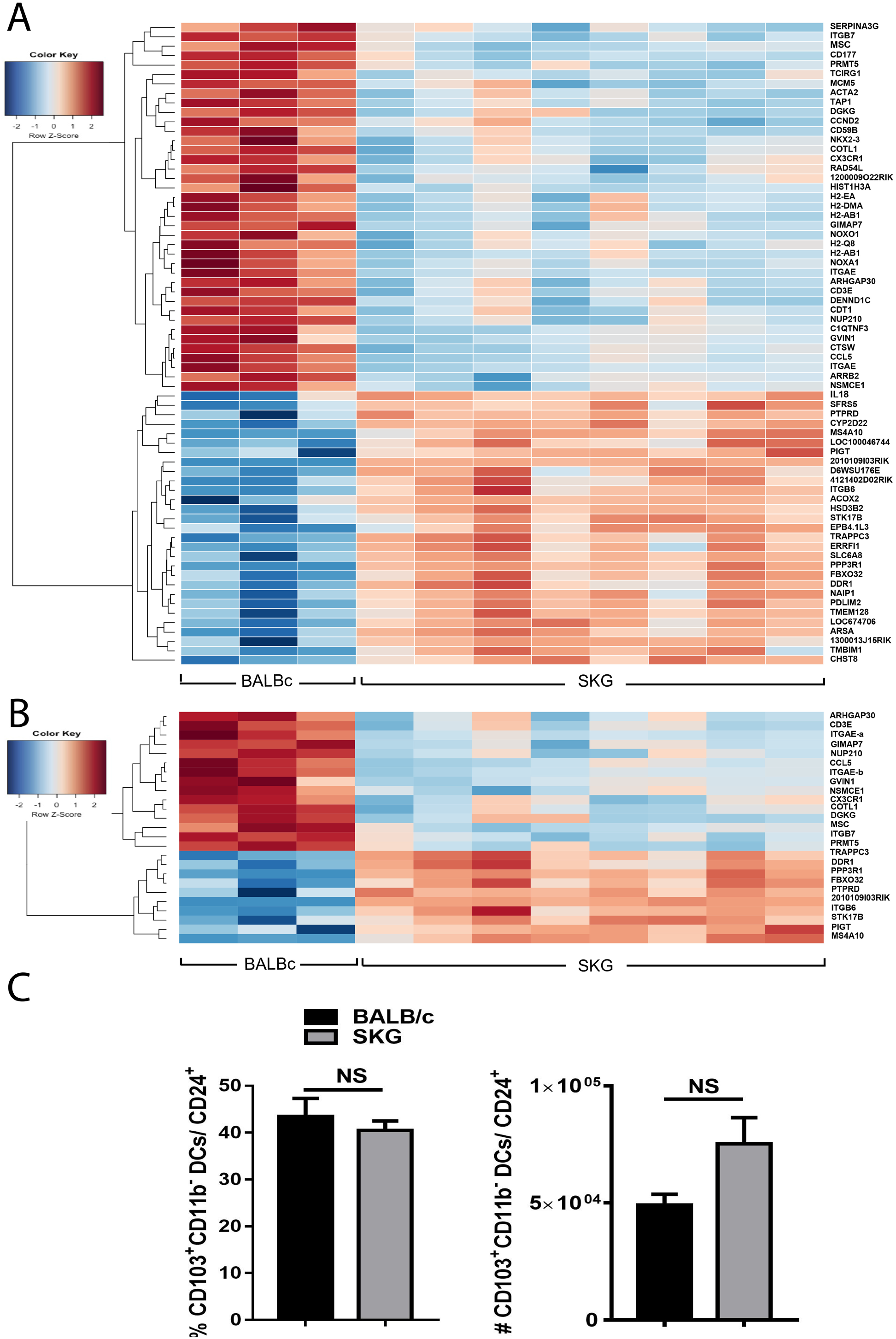
Differential expression of tissue resident memory T cell and Runx3-regulated genes in SKG ileum. **A**: Differentially expressed genes in ileum of BALB/c and SKG mice. The subset of Runx3-regulated genes is shown in (**B**). **C:** Percentage gated and total number of CD103^+^CD11b^-^ small intestine lamina propria DCs from BALB/c and SKG mice. Data are mean ± SEM. Representative of three separate experiments analyzing individual mice. n=3-5 per group. NS=not significant.

### The CD4^+^ T cell-specific SKG mutation impairs CD4-IEL cell trans-differentiation but not pTreg induction *in vitro* and *in vivo*

To assess the specific impact of the SKG CD4^+^ T cell ZAP70 mutation on CD4-IEL cell differentiation, we cultured ovalbumin (OVA)-specific naïve CD4^+^ T cells from DO11.10 (BALB/c mice with transgenic expression of OVA specific TCR) and SKG DO11.10 (SKG mice crossed to DO11.10) mice for 3 days with BALB/c DCs in presence of OVA_323-339_, TGF-β and/or RA, to induce CD8α and Foxp3 expression [12, 32]. Exogenous TGF-β alone or in combination with RA induced CD8α expression by DO11.10 CD4^+^CD25^-^ T cells but not SKG DO11.10 CD4^+^CD25^-^ T cells (Figure 3A). CD4^+^CD25^+^ Treg from BALB/c but not SKG mice also acquired CD8α expression (Figure 3B). In contrast, naïve T cells from both DO11.10 and SKG DO11.10 differentiated to Foxp3^+^ Treg in the presence of RA and TGFβ (Figure 3C). These data indicate that when signaled by BALB/c APCs, the ZAP-70^W163C^ mutant SKG TCR supports TGF-β and RA-mediated Foxp3^+^ pTreg but not CD4-IEL development. The data accord with the increased proportions of Treg and severe reduction in CD4-IEL in SI of SKG mice (Figure 1A). Consistent with this conclusion, proliferating BALB/c and SKG CD4^+^ T cells expressed comparable levels of TGF-β receptor 1 (Figure 3D and Supplementary Figure 3). Treg expansion in SKG relative to BALB/c mice may also be facilitated by increased proliferation, as indicated by increased Ki67^+^ cells among SKG T cells stimulated with or without TGFβ and RA in vitro (Supplementary Figure 3), and among SKG Foxp3^+^ Treg in vivo [33].

**Figure 3.**
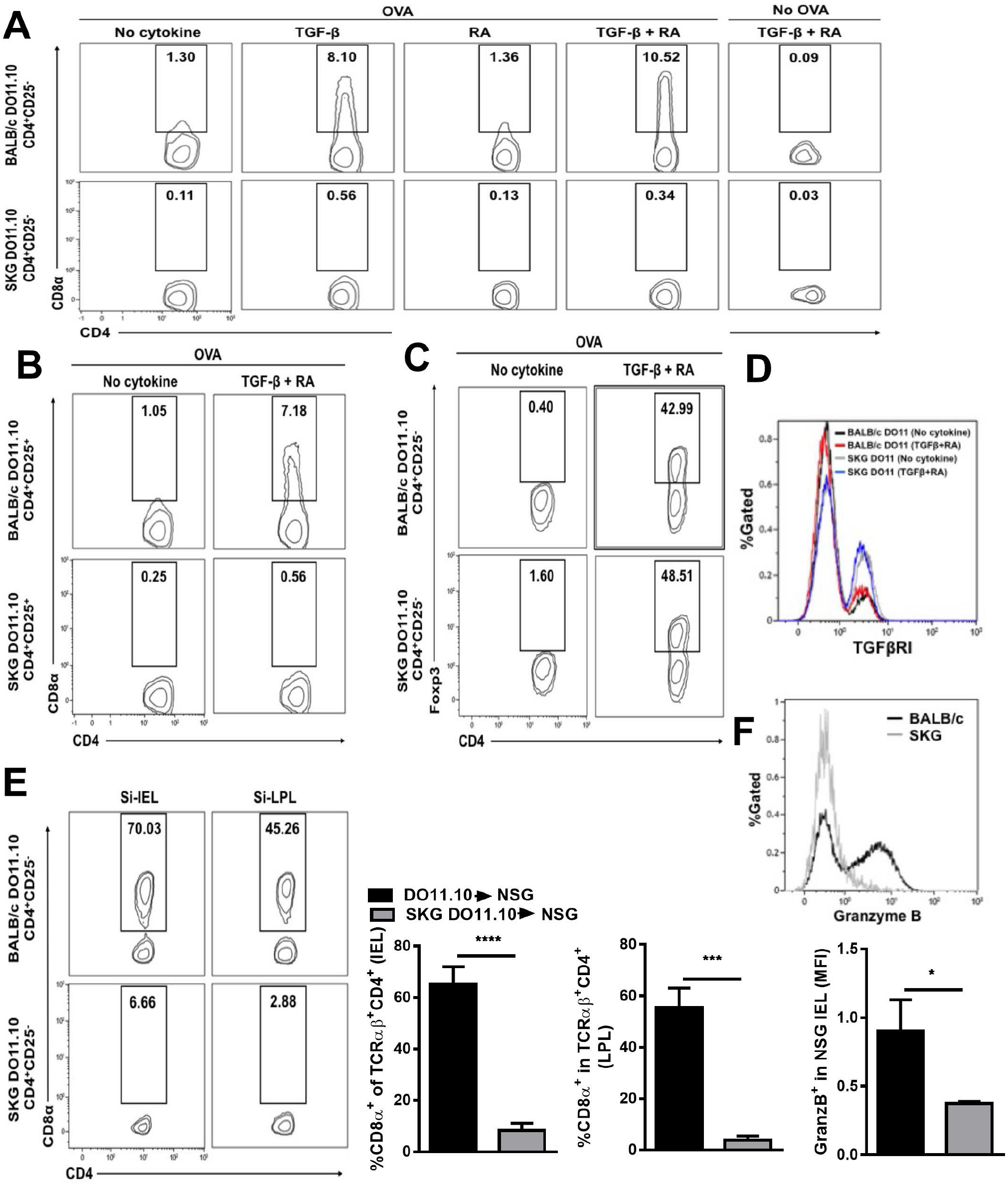
The CD4^+^ T cell intrinsic SKG mutation impairs CD4-IEL cell differentiation in vitro and in vivo. **A**: Expression of CD8α by sorted DO11.10 TCRβ^+^CD4^+^CD25^-^ T cells from BALB/c DO11.10 and SKG DO11.10 mice cultured with naïve BALB/c derived DCs stimulated with or without OVA peptide in the absence (no cytokine) or presence of TGF-β or RA or both. B: Expression of CD8α by DO11.10 TCRβ^+^CD4^+^CD25^+^ T cells isolated from BALB/c DO11.10 and SKG DO11.10 mice cultured with naïve BALB/c DCs and OVA peptide in the absence (no cytokine) or presence of TGF-β and RA. Expression of Foxp3 (**C**) and cell surface TGF-β receptor 1 (**D**) by BALB/c and SKG derived DO11.10 TCRβ^+^CD4^+^CD25^-^ T cells cultured as in **B. E, G**: Surface expression of CD8α by DO11.10 TCRβ^+^CD4^+^ T cells from the SI-IEL and SI-LPL of NOD SCID recipient mice 16 weeks after adoptive transfer of naïve TCRβ^+^CD4^+^CD8α^-^CD25^-^ splenic T cells purified from BALB/c DO11.10 and SKG DO11.10 mice. **F**: Intracellular staining of granzyme B in gated DO11.10 TCRβ^+^CD4^+^ T cells from the SI-IEL of NSG recipients. Representative of at least 2 separate experiments. Data are represented as mean ±SEM. * *p*<0.05, ** *p*<0.01, *** *p*<0.001, t test.

We therefore investigated whether CD4^+^ T cells from DO11.10 and SKG DO11.10 mice could transdifferentiate into CD4+CD8+ DP IEL *in vivo*. Sort-purified naïve CD4^+^CD25^-^ T cells from DO11.10 and SKG DO11.10 mice were adoptively transferred to immunodeficient NSG mice, to which BALB/c splenic DCs were transferred the day before, and fed with 1% OVA in the drinking water. Corroborating the *in vitro* results, significantly higher proportions of CD4^+^ T cells from DO11.10 than SKG DO11.10 mice transdifferentiated to DP, populating the IEL and LP environments 120 days post transfer (Figure 3E). SKG T cells also failed to upregulate cytotoxic activity as indicated by decreased expression of granzyme B (Figure 3F). Taken together these results indicate that CD4^+^ T cells from SKG mice recognize cognate antigen presented by DCs, sufficient to induce proliferation and TGF-β responsiveness but not trans-differentiation to DP cells.

### Inhibition of ZAP-70 kinase activity abrogates the development of CD4-IEL but not pTreg induction

Cognate antigen recognition is critical for cytokine-induced CD4^+^ T cell differentiation in the periphery. The SKG mutation, although not located in the catalytic site of Zap70, decreases TCR activation-induced PKCθ phosphorylation and impairs downstream TCR signaling in T helper cells [23]. To determine whether Zap70 kinase activity regulates CD4-IEL development, we cultured naïve CD4^+^CD25^-^ or CD4^+^CD25^+^ T cells from DO11.10 mice with OVA-pulsed splenic DCs in presence of TGF-β and RA, and increasing doses of ZAP-70 kinase inhibitor, piceatannol. Piceatannol dose-dependently inhibited the development of DP cells (Figure 4A) but did not impair differentiation to Foxp3^+^ pTreg (Figure 4B). CD4^+^CD25^-^ and CD4^+^CD25^+^ T cell Runx3 and CD103 expression induced by TGF-β and RA were also inhibited in the presence of piceatannol (Figure 4C, D). In contrast, T cell activation as measured by CD69 expression was barely affected by piceatannol (Figure 4E). These findings indicate that TCR activation-induced Zap70 kinase activity differentially regulates TGF-β and RA-induced pTreg induction and transcriptional reprogramming of CD4^+^ T cells to trans-differentiate into DP cells.

**Figure 4.**
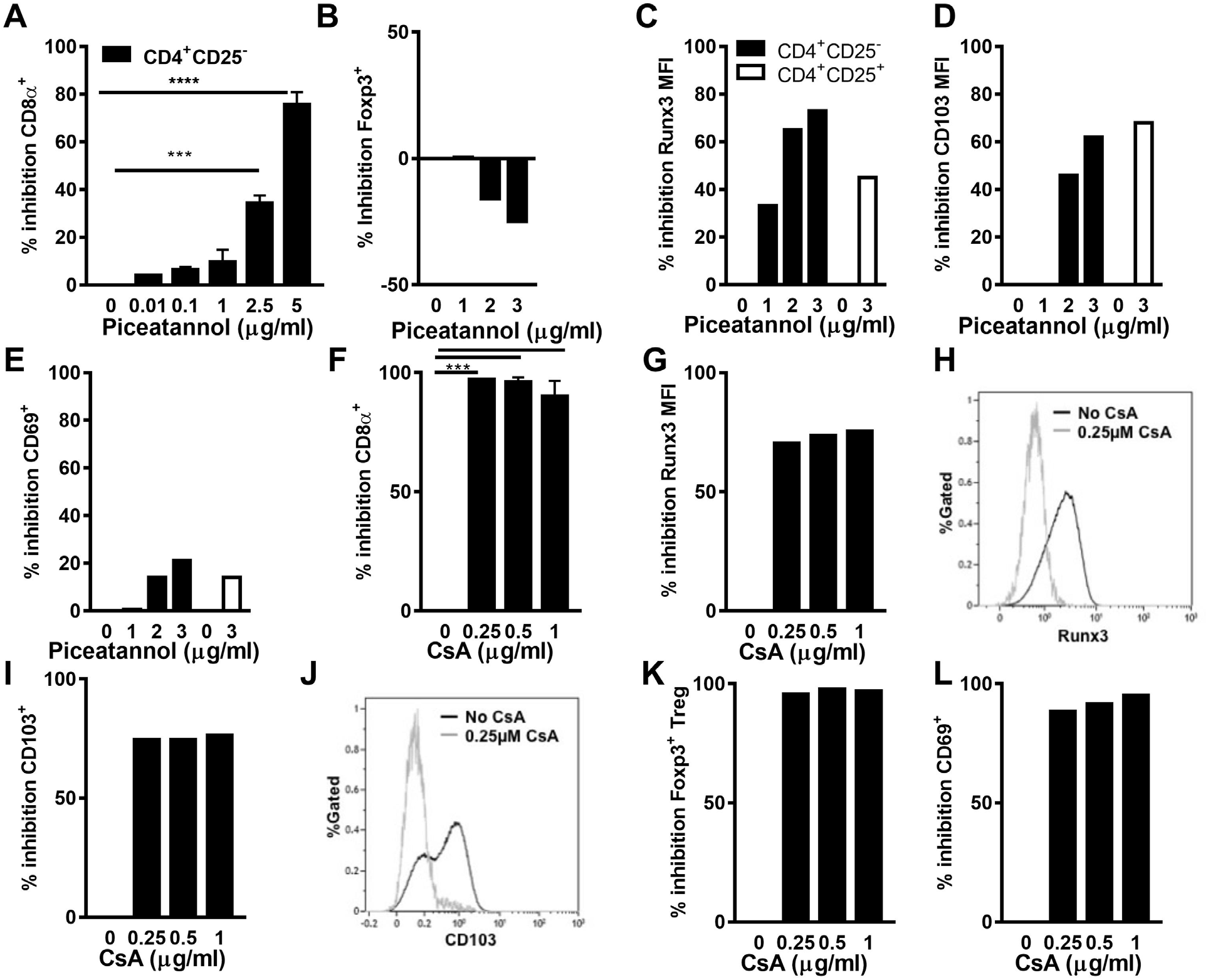
Inhibition of ZAP-70 kinase activity and calcium signaling suppresses CD4 T cell differentiation to CD4-IEL. Inhibition of CD8α (**A**) or Foxp3 (**B**) expression by sorted DO11.10 TCRβ^+^CD4^+^CD25^-^ splenic T cells cultured with OVA-pulsed DCs in presence of TGF-β, RA and increasing concentrations of Piceatannol. Inhibition of Runx3 (**C**), CD103 (**D**), and CD69 (**E**) expression by sortedDO11.10 TCRβ^+^CD4^+^CD25^-^ and DO11.10 TCRβ^+^CD4^+^CD25^+^ T cells cultured with OVA-pulsed DCs in presence of TGF-β, RA and increasing concentrations of Piceatannol. Inhibition of CD8α (**F**), Runx3 (**G, H**), CD103 (**I, J**) and Foxp3 (**K, L**) by purified BALB/c derived DO11.10 TCRβ^+^CD4^+^CD25^-^ T cells cultured with OVA pulsed DCs in presence of TGFβ, RA and increasing concentrations of cyclosporine A (CsA).

### Calcium signaling is required for TGF-β and RA-mediated T helper cell trans-differentiation to CD4-IEL and pTreg

TCR activation and downstream ZAP-70 signaling result in calcium influx, which is reduced upon TCR activation in CD4^+^ T cells from SKG mice [23]. Therefore, we cultured DO11.10 naïve CD4^+^ T cells under CD4-IEL trans-differentiating conditions in the presence of 0.25 µM cyclosporine A (CsA), which inhibits calcineurin-mediated calcium influx. CsA prevented cytokine-induced CD4^+^CD25^-^ T cell trans-differentiation to CD4-IEL in the presence of OVA_323-339_ (Figure 4F) and inhibited upregulation of Runx3 (Figure 4G, H) and CD103 (Figure 4I, J). In contrast to piceatannol, pTreg induction (Figure 4K) and CD69 expression (Figure 4L) were inhibited by CsA treatment, indicating that inhibition of calcium signaling suppresses TCR activation and abrogates the TGF-β and RA-induced CD4^+^ T cell differentiation program to CD4-IEL and Treg *in vitro*.

### CD4-IEL are reduced in AS patients

SKG mice develop the clinical syndrome of SpA with curdlan administration. While naïve SKG mice appear healthy, they have gut dysbiosis and intestinal immune dysregulation [24]. *ZAP70* variants are not associated with human SpA risk, but AS patients have similar intestinal dysbiosis and immune dysregulation [3, 34]. To identify potential similarities in signalling pathways, we compared differentially expressed genes in SKG relative to BALB/c ileum, with risk SNP identified in GWAS studies of AS, IBD, Primary Sclerosing Cholangitis and Psoriasis risk [35, 36]. *NKX2-3* and *TMBIM1* are associated with AS, Crohn’s disease, Ulcerative Colitis and Primary Sclerosing Cholangitis, *ERRFI1* with Ulcerative Colitis and *HLA-DQB1* and *HLA-DQB2* with IBD (Figure 5A). Consistent with its downregulated expression in SKG ileum, the promoter of *NKX2-3*, which regulates intestinal lymphocyte homing in mice, binds the downstream signalling molecule NFAT1 less efficiently than the protective variant [37]. To more directly examine this, we explored CD4-IEL frequency in ileal biopsies of a small number of AS patients and healthy controls (HC). Previously we demonstrated that Foxp3^+^ Treg were expanded in AS relative to HC intestine [34]. Similar to SKG mice, the proportion of CD4-IEL was reduced among ileal T cells in AS compared with HC (Figure 5B). Among PBMC, %DP CD3^+^TCRγδ^-^TRAV7.2^-^CD56^-^ T cells was significantly reduced and %CD25^hi^CD127^-^ Treg significantly increased in patients with AS compared with HC. %CD8^+^ T cells was also decreased and %monocytes and B cells increased in AS patients relative to HC (Figure 5C). In a second cohort of AS patients treated with or without TNFi, the proportion of CD3^+^ DP that was CD4^lo^CD8^hi^ was significantly reduced relative to HC in AS patients not taking TNFi, but not in AS patients taking TNFi (Figure 5D). CD4^lo^CD8^hi^ DP increased in each of 4 AS patients after commencing treatment with TNFi. These data suggest that similar to SKG mice, CD4-IEL differentiation is impaired in AS patient SI, and the ratio of DP:Treg is reduced. The increase in relative proportion of CD4^lo^CD8^hi^ DP in AS patients commencing TNFi further suggests that impaired CD8 expression may be correctible in AS CD4^+^ T cells.

**Figure 5.**
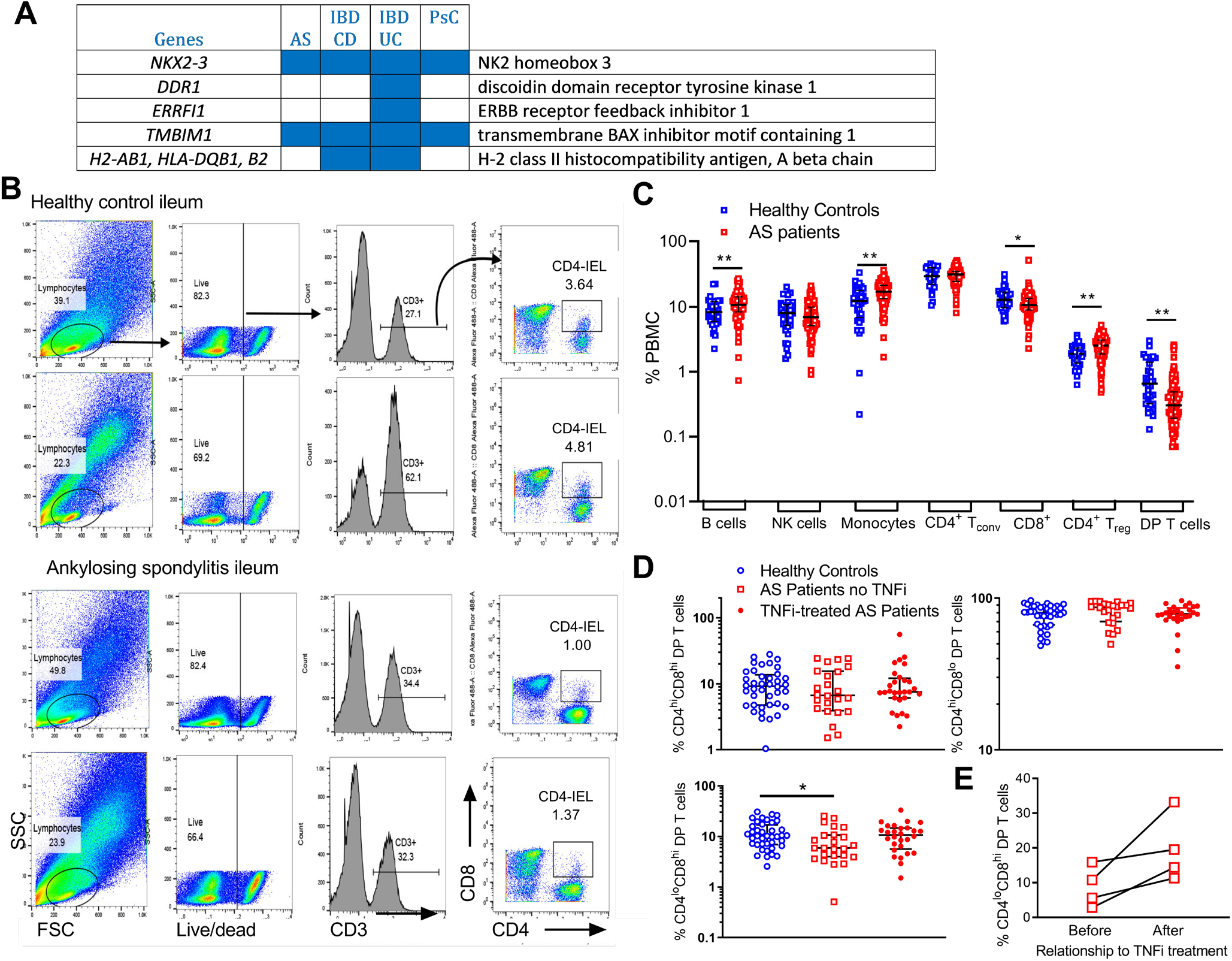
Intestinal CD4-IEL and circulating double positive T cells are reduced in patients with Ankylosing Spondylitis. (**A**) DE genes in SKG relative to BALB/c ileum, in common with risk SNP identified in GWAS studies of AS, Primary Sclerosing Cholangitis (PsC) and Psoriasis risk [36] and IBD risk [35]. (**B**) Flow cytometry plots showing gating strategy and frequency of CD4-IEL from ileum of two patients with active AS and two healthy controls. (C) Analysis of PB B cells, NK cells, classical monocytes, CD8+ T cells, conventional CD4+ (Tconv), regulatory CD4+ (Treg) and double positive (DP) T cells from 128 AS patients and 34 healthy controls (cohort 1) based on gating strategy in Supplementary Figure 4A. All data are shown with median and IQR. * *p*<0.05, ** *p*<0.01 for each cell population in AS patients compared to controls (Mann-Whitney U test). (D) Analysis of PB DP T cells from 24 untreated AS patients, 28 AS patients treated with TNF inhibitors and 40 healthy controls (cohort 2), based on gating strategy in Supplementary Figure 4B. All data are shown with median and IQR. * *p*<0.05 for untreated AS compared to controls for the CD4^lo^CD8^hi^ population (Kruskal-Wallis test with Dunn’s correction for multiple testing). (E) Analysis of PB CD4^lo^CD8^hi^ DP T cells from 4 AS patients before and after commencing treatment with TNFi. *p*=0.058 for comparison before and after TNFi (ratio paired t test).

## Discussion

The results presented in this study extend the understanding of mature CD4^+^ T lymphocyte differentiation in the intestine by demonstrating that TCR signaling strength influences TGF-β and RA-mediated acquisition of regulatory and cytotoxic phenotypes. W163C is a missense mutation located in the C-terminal domain of Zap70, which decreases the interaction of Zap70 with TCR-ζ and reduces TCR signal strength [23]. We show that this weak TCR activation specifically prevents TGF-β and RA-induced post-thymic acquisition of CD8α, but not Foxp3, by mature CD4^+^ T cells, indicating that high affinity TCR signaling combined with TGF-β and RA signaling promotes CD4-IEL cell development in the intestinal epithelium. While TGF-β and RA derived from CD103^+^ DCs also induce Foxp3^+^ Treg in the gut [11], we show here that Foxp3^+^ Treg differentiation was not affected by the reduced TCR signal strength due to Zap70^W163C^. Only a subset of antigen-specific CD4^+^ T cells from wt mice converts to CD4-IEL *in vitro* and it is possible that only these cells recognize antigen with sufficiently high affinity for trans-differentiation. Consistent with a requirement for high affinity TCR activation in vivo, intestinal CD4-IEL demonstrate a highly activated phenotype [12] and develop from epithelial-infiltrating CD4^+^ T cells or Foxp3^+^ Treg [14], which may have been selected by agonist antigens in the thymus [13].

Although weak TCR signaling may impair thymic Treg cell development and thymic T cell output [23], our results show that Zap70^W163C^ does not inhibit and indeed favors intestinal pTreg induction in SKG mice, notwithstanding that CD4 lymphopenia constrains pTreg numbers. The reduced thymic T cell output in SKG mice and the negligible levels of CD4-IEL in SKG intestine despite increased pTreg indicate that the mature SKG CD4^+^ T cell and Treg precursors of CD4-IEL lack the capacity to transduce strong antigen stimulation upon migration to intestinal epithelium. Similar to SKG mice, we showed previously that TCR diversity is increased, T cell expansions reduced and Epstein-Barr virus-specific clonotypes expanded in the AS relative to healthy control TCR repertoire, consistent with reduced capacity for TCR signaling by strong antigens and reduced control of latent viral infection [25]. Although only a small number of intestinal samples was obtained, these data support the reduction in CD4-IEL that we observed in AS intestine.

Mutation-associated defects in Zap70 function or expression lead to specific defects in the thymic development and survival of CD8^+^ but not CD4^+^ T cells in SKG and other mouse models [23, 38, 39]. People with mutations in Zap70 similarly develop severe combined immunodeficiency due to deficient CD8^+^ T cell development and production of CD4^+^ T cells with deficient TCR signaling [40, 41]. In these patients, loss of Zap70 is compensated by splenic tyrosine kinase (Syk), which is overexpressed in thymocytes at later stages of development, and promotes CD4^+^ T cell development and TCR responsiveness of residual CD8^+^ T cells [42]. In DCs, dectin-1 signaling by β-glucan was shown to activate syk and raf-1 protein tyrosine kinases to trigger NF-κB and cytokine production [43]. Compensatory upregulation of Syk may facilitate CD4^+^ T cell development and their differentiation to Treg and Th17 cells in the periphery of SKG mice, particularly after delivery of β-glucan [43]. However, TGF-β and RA stimulation failed to stimulate SKG CD4^+^CD25^-^ T cells or Treg to trans-differentiate to CD4-IEL, curbing expression of CD8, granzyme B and Runx3. Furthermore, chemical inhibition of Zap70 produced comparable defects in BALB/c CD4 T cells. Genetic or chemical disruption of Zap70 catalytic activity demonstrated that Zap70 is required for immune synapse formation between CTL cells and their targets, as well as cytotoxicity and cytokine production [44]. These observations support an indispensable role for Zap70-mediated TCR signaling in the development and function of CD8^+^ CTL and IEL.

Besides CD4-IEL trans-differentiation in the intestine, TGF-β-induced Runx3 is also required for Foxp3 expression [45-47]. Thus, Runx3 plays dual roles in the control of intestinal regulation. *RUNX3* variants are associated with AS and psoriatic arthritis SpA risk [1]. A putative regulatory element upstream of *RUNX3* containing the risk SNP significantly reduced expression of *RUNX3* and recruitment of IRF4 to the nucleus relative to the protective allele [2]. In CD8^+^ T cells, an AS-risk allele of *RUNX3* was shown to bind components of the NuRD complex and Aiolos more strongly than the protective allele [21]. Our data demonstrating lower levels of DP T cells in SI and blood in AS are consistent with this deficiency, and increases in CD4^lo^CD8^hi^ DP with TNF inhibition implicate epigenetic regulation, potentially through the NuRD complex [48], which is central to repression of Runx3 gene expression. Furthermore, Runx3 expression was reduced in CD4^+^ T cells in SKG mice and Runx3-regulated genes account for a high proportion of the DE genes in SKG ileum. Together our data link Runx3 with CD4-IEL deficiency, intestinal pathology and AS risk.

In mice, gut microbiota and indoles control intestinal CD4-IEL development [22]. In healthy human colon, some CD4-IEL responded strongly to the commensal bacterium, *Faecalibacterium prausnitzii* and IEL were reduced in colonic mucosa of IBD patients. *F. prausnitzii* deficiency is associated with human IBD severity [49, 50]. Several lines of evidence indicate that CD4-IEL confer protection against spontaneous colonic inflammatory diseases [13, 31], including compensation for loss of Treg and prevention of inflammatory responses [13]. Taking together the similar features in SKG mice and AS, our data indicate that deficiency of CD4-IEL relative to Treg is a common feature of SpA. Furthermore, CD4^+^ T cell specific TCR Zap70 activity acts as a key “switch” between Treg conversion and CD4-IEL trans-differentiation, and Runx3 deficiency is a common link between CD4^+^ T cell function in mouse and human SpA.

## Contributorship

Study concept and design: ZAB, RT. Acquisition and analysis and interpretation of data: all authors. Drafting of the manuscript: ZAB, RT. Critical revision of the manuscript for important intellectual content: all authors. Obtained funding: RT.

## Acknowledgements

We thank James McCahon, and the TRI biological research and flow cytometry core facilities for assistance with experiments.

## Ethical approval information

Studies were approved by the Human Research Ethics Committee of the Universitá della Campania teaching hospital (approval no. Prot. 0013336/i), the Metro South (approval no. HREC/05/QPAH/221) and the Queensland University of Technology (QUT) Human Research Ethics Committees (approval no. 1600000162).

## Public and patient involvement

At what stage in the research process were patients/the public first involved in the research and how? Patients and the public were involved at the recruitment stage.

How were the research question(s) and outcome measures developed and informed by their priorities, experience, and preferences? No direct patient involvement.

How were patients/the public involved in the design of this study? Not involved.

How were they involved in the recruitment to and conduct of the study? Not involved.

Were they asked to assess the burden of the intervention and time required to participate in the research? No.

How were (or will) they be involved in your plans to disseminate the study results to participants and relevant wider patient communities (e.g. by choosing what information/results to share, when, and in what format)? Results will be shared via a newsletter to the involved participants to inform them of study results.

