## Supplemental material for "Deficiencies of Runx3 and tissue-resident CD4^+^ intestinal epithelial lymphocytes link intestinal dysbiosis and inflammation in mouse and human spondyloarthropathy"

**Supplementary material**

**Supplementary methods**

**Mice**

SKG mice, originally obtained from S. Sakaguchi (University of Kyoto), DO11.10 and NSG mice were bred and housed under SPF conditions at The University of Queensland Translational Research Institute Animal Facility. Approval for all experiments was obtained from The University of Queensland animal ethics committee. BALB/c were purchased from the Animal Resources Centre. SKG DO11.10 mice were generated by crossing SKG and DO11.10 mice. Male and female mice (n=12-15 per group including repeats, up to 60 per experiment, 1200 for entire study) were used between 8-12 weeks of age. All analyses carried out by an investigator blinded to group. No data were excluded from the analysis or figures. Group size based on initial experiments demonstrating effect size and variance of parameters studied. Litters were randomised over the groups and were age-matched to availability. Treatment order and measurement by cage was randomised.

**Isolation of intraepithelial and lamina propria lymphocytes**

Small intestines were collected and Peyer’s patches removed. Intestines were cut open longitudinally and intestinal luminal contents were washed with cold PBS, then cut into 1 cm pieces. The pieces were stirred at 37 °C for 20 min in HBSS supplemented with 5% FCS, penicillin/streptomycin and 3 mM EDTA. The intraepithelial lymphocytes (IELs) in the supernatant were collected and washed. Residual tissue was cut into fine pieces and stirred in RPMI 1640 supplemented with 10% FCS, collagenase D (Roche) and DNase I (Sigma-Aldrich) at 37 °C for 40 min. The lamina propria lymphocytes (LPLs) in the supernatant were collected and washed with RPMI 1640.

**Antibodies and flow cytometry**

Fluorochrome-conjugated mAb specific for mouse CD4, CD45.2, CD8a, CD8b, CD103, CD25, CD44, CD69, TCRb, TCRgd, DO11.10 TCR were purchased from Biolegend. Antibodies specific for ThPOK and Runx3 were purchased from BD Biosciences. Foxp3 mouse regulatory kit (ebioscience) was used for intracellular staining of Foxp3. Cell populations were first stained with cell surface marker-specific mAb, then permeabilized in Fix/Perm buffer and stained in Perm/wash buffer. Fluorochrome-conjugated mAb specific for human markers are detailed in Supplementary Table 3. Flow cytometry data were acquired on a Gallios cytometer (Beckman Coulter) and were analysed with Kaluza V1.3 software. Gating strategy for PBMC from the UK cohort is shown in Supplementary Figure 4A and for PBMC for the Australian cohort in Supplementary Figure 4B.

**In vitro T cell culture**

Naïve (CD4^+^CD25^-^) T cells were isolated from DO11.10 and SKG DO11.10 mice using the CD4^+^ T cells isolation kit (Miltenyl Biotec) and were cultured with CD11c^+^ splenic DC in the presence of 500 nM OVA peptide, TGF-β (2 ng/ml, R&D Systems) and RA (10 nM). In some experiments, Piecetannol (Sigma-Aldrich) or cyclosporine A (Sigma-Aldrich) were added.

**In vivo transfer experiment**

FACS purified 5x10^5^ naïve (CD4+CD45RBhiCD25-) T cells from DO11.10 or SKG DO11.10 mice (5 mice pooled for each transfer) and 1x10^6^ MACS purified splenic CD11c+ DC from BALB/c mice were transferred to recipient NSG mice. NSG mice were fed with 1% chicken OVA (Sigma-Aldrich) after T cell transfer. At 4-16 weeks after transfer, donor cells were isolated from intestine of the NSG recipient mice and analyzed by flow cytometry.

**Microarray**

Total RNA was purified from terminal ileum of 8 naïve SKG and 3 naïve BALB/c mice using RNeasy kits (Qiagen). RNA from tissues selected for microarray (10 - 30mg, Bioanalyzer RNA Integrity Scores >6) was amplified using the TotalPrep RNA amplification kit (Ambion). Amplified RNA was hybrized to Illumina MouseRef-8 v2.0 Expression BeadChip arrays. Arrays were scanned on an Illumina iScan.

**Patients**

Two to four adjacent mucosal biopsies were obtained from the ileum of each patient and each control. Control blood samples were sourced from unrelated healthy white subjects with no spondyloarthritis symptoms. Written informed consent was received from all participants.

**Single cell suspension and staining of intestinal biopsies**

Lamina propria mononuclear cells (LPMC) were isolated from the gut of patients with AS, and healthy controls as previously described. In brief, cryopreserved small intestinal biopsies were rapidly thawed in a 37°C water bath and washed in 20 mL R10 before tissue dissociation. Ileal samples were incubated in R10 media with 1 mg/ml Collagenase D (Roche) and 100 mg/ml DNase (Thermo Fisher Scientific) for one hour. Biopsies were then dissociated by vigorous agitation using a GentleMACS Dissociator (Miltenyi Biotec), then strained through a 70 mm filter. Cells were washed with R10 media and collected for subsequent flow cytometry analysis: after washing cells twice using FACS buffer (Phosphate-buffered saline (PBS**)** with the addition of 1% Fetal Bovine Serum (FBS)).

**Flow cytometry**

Staining buffer was prepared by adding flourochrome-conjugated antibodies and fixable dyes to FACS buffer. Monoclonal antibodies from Biolegend (San Diego, California) included: anti-CD3-Brilliant Violet 421 (clone SK7), anti-CD4-peridinin chlorophyll protein (PerCP)-Cy5.5 (clone RPA-T4), anti-CD8-FITC (clone RPA-T8), anti-CD3-PE (OKT3), anti-CD8-APC-Cy7 (HIT8a) and anti-CD4-APC (RPA-T4). Live/dead staining was performed with the eFluor 780 Fixable Viability Dye (Thermofisher). Cells were then incubated for 20 minutes at 4°C in the dark. After staining, cells were washed twice with 200 μl FACS buffer and resuspended in 200 μl fixing buffer (PBS with the addition of 3% paraformaldehyde) before acquisition on a FACSCanto II Flow Cytometer (gut, BD Biosciences) or MoFlo Astrios (PBMC, Beckman Coulter) with the use of the FlowJo software (BD Biosciences). Cells were expressed as percentage of cells within the lymphocyte gate. Isotype control values were subtracted in all the samples analysed.

**Statistics**

Except for the microarray, Graph Pad Prism software was used for statistical analysis. Murine data were analysed by one-way ANOVA or unpaired student’s t-test as indicated. Human data, having tested for skewness, were analysed by Mann-Whitney test, Kruskal-Wallis test with Dunn’s correction for multiple testing and paired t test as indicated.

Microarray data were initially extracted through Illumina GenomeStudio. We removed batch effects using the limma package in R. Raw data were imported into Genespring and normalized to the 50th percentile (per array) and each gene was normalized to its median expression across all arrays. Probes with Illumina detection values ≥0.95 in all arrays were retained for analyses. Gene expression data have been deposited in the Gene Expression Omnibus (GEO) database GSE106134. Genes differentially expressed in SKG mice relative to BALB/c were uncovered by fitting a linear model (lmFit function in limma package). We used eBayes moderation to adjust the standard error for gene models. We adjusted p-values (padj) using False Discovery Rate (FDR) multiple correction method. The differentially expressed genes with padj ≤ 0.05 are shown.

We searched RUNX3-regulated genes in CD8^+^ T cells uncovered using ChIP-seq and transcriptome analysis (GEO accession number GSE50131, [29]), for enrichment in SKG and BALB/c mice. Disease targets of differentially expressed RUNX3-modulated genes were identified using Open Targets.

**Supplementary Figure S1. Helios expression by Foxp3+ Treg in LPL**
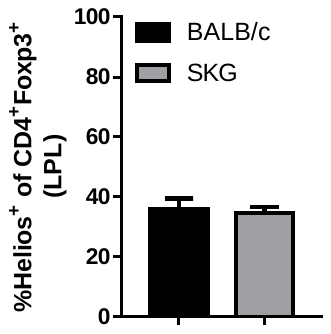

**Supplementary Figure 2. CD103^+^CD11b^-^ DCs in the intestinal lamina propria of BALB/c and SKG mice**

(**A**) Gating strategy to identify retinoic acid producing CD103^+^CD11b^-^ lamina propria DCs in naïve BALB/c mice. (**B**) Mean fluorescence intensity of surface MHC class-II expression by CD103^+^CD11b^-^ dendritic cells. Data are represented as mean ±SEM. Representative of two separate experiments analyzing individual mice. n=5/group. **p<0.01 by Mann Whitney t- test.

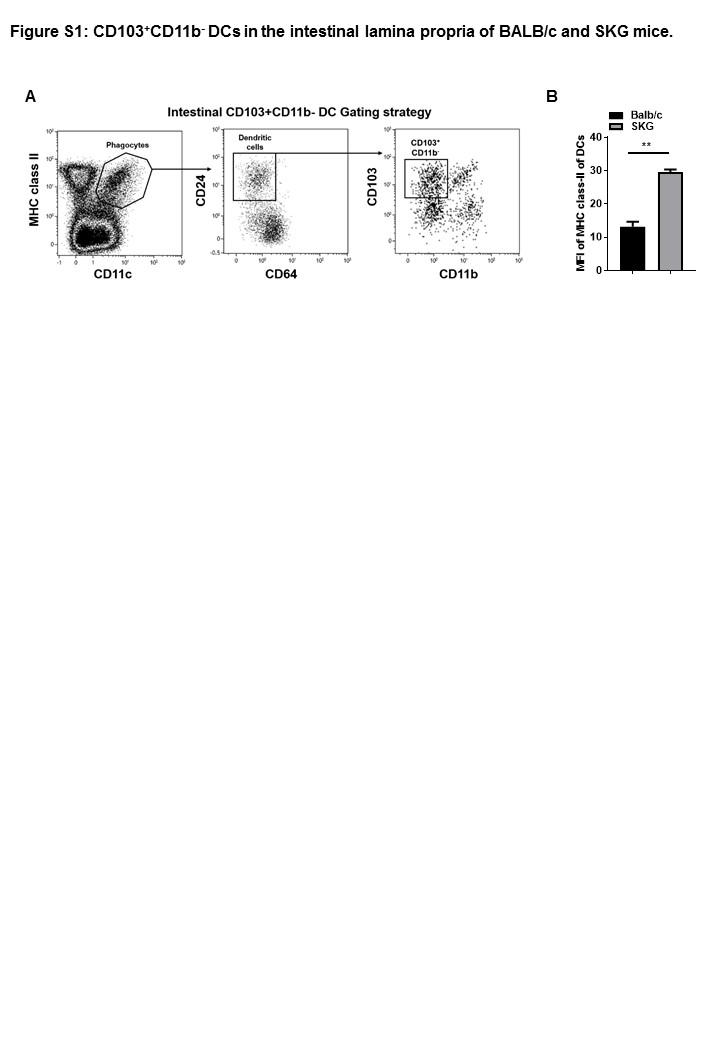

**Supplementary Figure 3. Proliferation of DO11.10 and SKG DO11.10 CD4^+^CD25^-^ cells in vitro**

(**A**) Staining of intracellular Ki67 and TGF-β receptor I by BALB/c and SKG derived DO11.10 TCRβ^+^CD4^+^CD25^-^ T cells cultured with DCs pulsed with OVA peptide in the absence (no cytokine) or presence of TGF-β or RA or both.

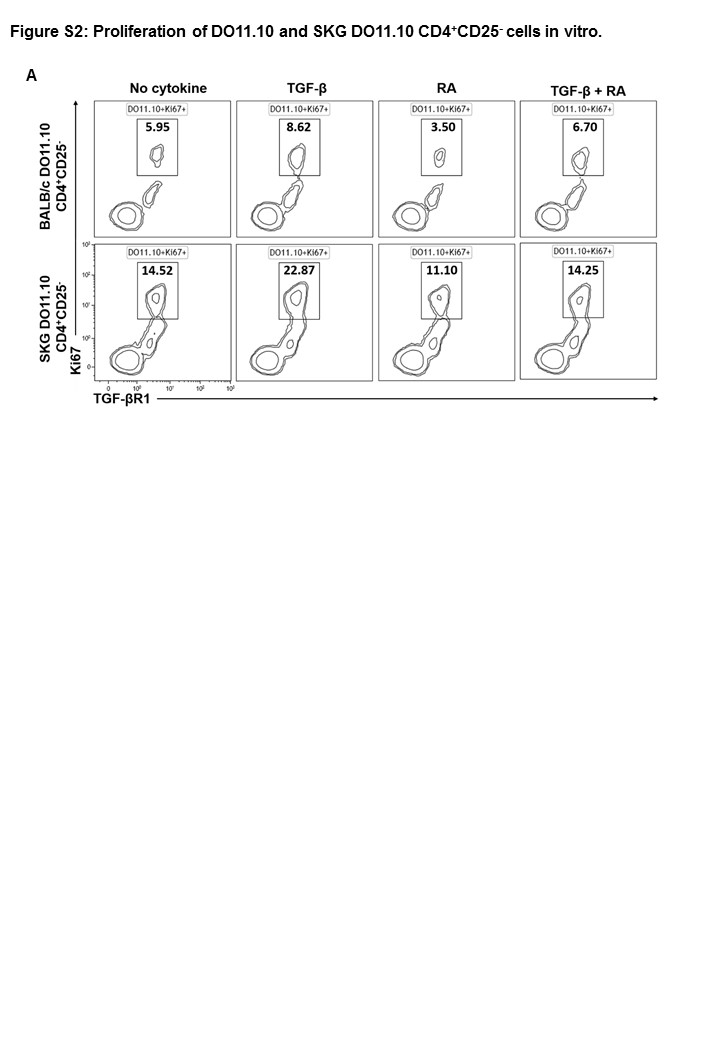

**Supplementary Figure 4. Gating strategies for PBMC.**

**A:** UK cohort. Relationships across plots are indicated by arrows. TCRαβ^+^ T cells were gated as CD19^+^CD3^+^CD56^-^γδTCR^-^Vα7.2^-^ in order to exclude NKT cells, γδ T cells and MAIT cells. After identifying CD8^+^ and CD4^+^ single positive T cells and CD4/8 DP populations, CD4^+^ Treg were gated as CD25^hi^CD127^lo^ or the residual conventional T cells (Tconv). Classical monocytes were gated as CD14^+^CD16^-^, B cells as CD19^+^ and NK cells as CD56^+^.

B: Australian cohort. Live CD3^+^CD4/8 DP T cells were identified and subdivided into CD4^hi^CD8^lo^, CD4^hi^CD8^hi^ and CD8^hi^CD4^lo^ populations.

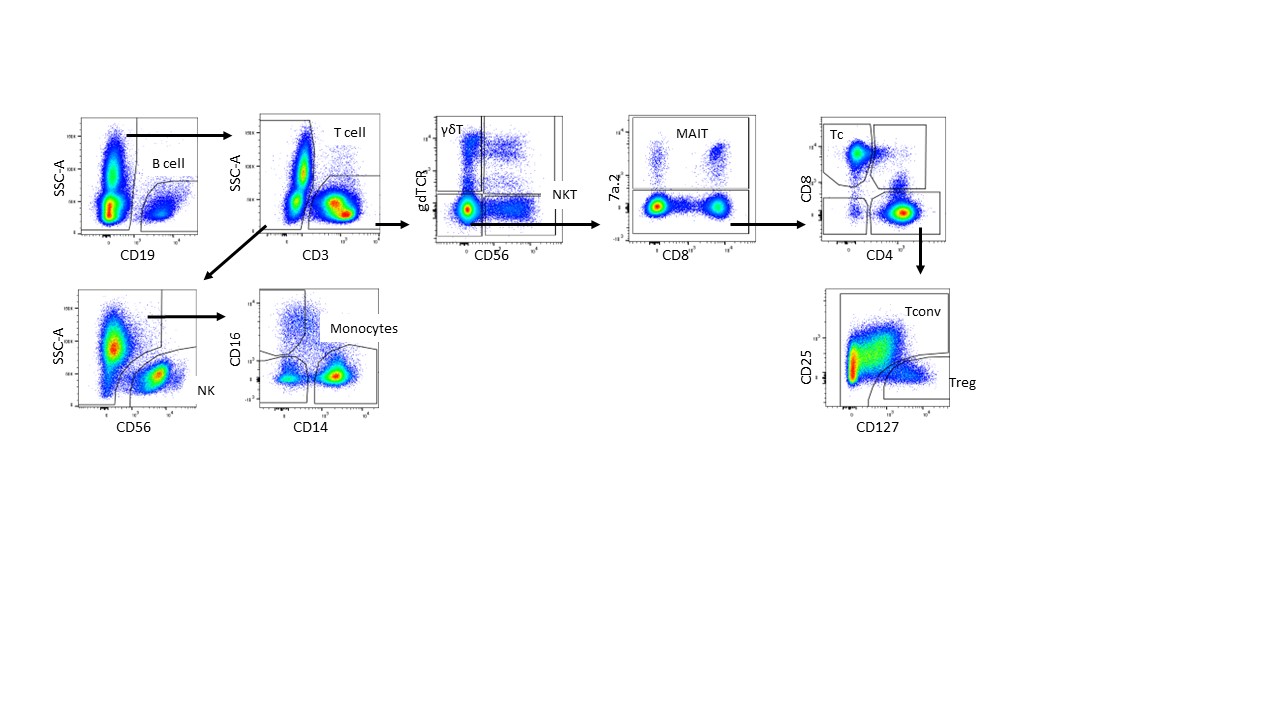
**A**

**B**

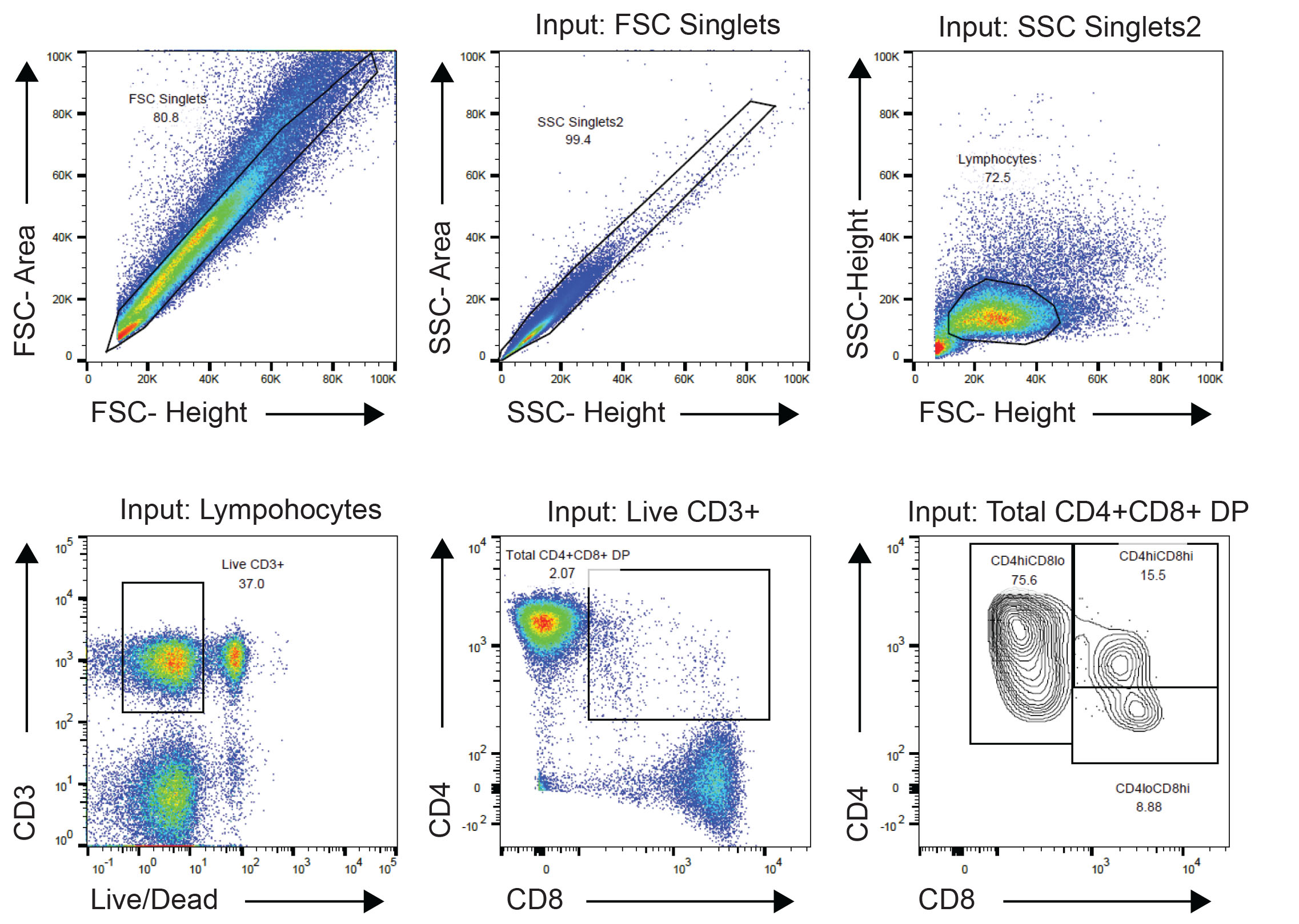

**Supplementary Table 1. Gene Ontogeny Analysis for differentially expressed genes in SKG vs BALB/c ileum.**

| **GO.Term** | **Description** | **P value** | **FDR.q.value** | **DE genes** |
| --- | --- | --- | --- | --- |
| <GO:0048002> | antigen processing and presentation of peptide antigen | 2.45e-05 | 0.36 | 4 |
| <GO:0050870> | positive regulation of T cell activation | 2.9e-05 | 0.213 | 6 |
| <GO:1903039> | positive regulation of leukocyte cell-cell adhesion | 3.93e-05 | 0.192 | 6 |
| <GO:2000504> | positive regulation of blood vessel remodeling | 8.01e-05 | 0.294 | 2 |
| <GO:0022409> | positive regulation of cell-cell adhesion | 8.87e-05 | 0.261 | 6 |
| <GO:0042605> | peptide antigen binding | 0.000132 | 0.577 | 3 |
| <GO:0008305> | integrin complex | 0.000147 | 0.264 | 3 |
| <GO:0050776> | regulation of immune response | 0.000154 | 0.377 | 9 |
| <GO:0019882> | antigen processing and presentation | 0.00016 | 0.336 | 4 |
| <GO:0044459> | plasma membrane part | 0.000178 | 0.16 | 19 |
| <GO:0051251> | positive regulation of lymphocyte activation | 0.000183 | 0.336 | 6 |
| <GO:0098636> | protein complex involved in cell adhesion | 2e-04 | 0.12 | 3 |
| <GO:0042129> | regulation of T cell proliferation | 0.000208 | 0.34 | 5 |
| <GO:0016176> | superoxide-generating NADPH oxidase activator activity | 0.000278 | 0.609 | 2 |
| <GO:0050863> | regulation of T cell activation | 0.000308 | 0.453 | 6 |
| <GO:0042102> | positive regulation of T cell proliferation | 0.000354 | 0.474 | 4 |
| <GO:0060312> | regulation of blood vessel remodeling | 0.00037 | 0.453 | 2 |
| <GO:0002696> | positive regulation of leukocyte activation | 0.00038 | 0.43 | 6 |
| <GO:0016020> | membrane | 0.000409 | 0.183 | 40 |
| <GO:1903037> | regulation of leukocyte cell-cell adhesion | 0.000412 | 0.433 | 6 |
| <GO:0002250> | adaptive immune response | 0.00043 | 0.421 | 5 |
| <GO:0043020> | NADPH oxidase complex | 0.000475 | 0.17 | 2 |
| <GO:0042613> | MHC class II protein complex | 0.000475 | 0.142 | 2 |
| <GO:0050867> | positive regulation of cell activation | 0.000493 | 0.452 | 6 |
| <GO:0043235> | receptor complex | 0.000677 | 0.174 | 6 |
| <GO:0003823> | antigen binding | 0.000692 | 1 | 3 |
| <GO:0002697> | regulation of immune effector process | 0.000741 | 0.64 | 6 |
| <GO:0050670> | regulation of lymphocyte proliferation | 0.00076 | 0.621 | 5 |
| <GO:0032944> | regulation of mononuclear cell proliferation | 0.000796 | 0.616 | 5 |
| <GO:0098797> | plasma membrane protein complex | 0.000871 | 0.195 | 5 |
| <GO:0070663> | regulation of leukocyte proliferation | 0.000931 | 0.684 | 5 |

Supplementary Table 2. **RUNX3-regulated genes in CD8+ T cells uncovered using ChIP-seq and transcriptome analysis** **differentially expressed** **in SKG relative to BALB/c ileum.** Log fold change (FC), adjusted p-values (p_adj_) and gene description are shown. ChIP-seq and transcriptome analysis GEO accession number GSE50131. Genes shown twice are isoforms.

| Gene | logFC | *p*_adj_ | Gene description |
| --- | --- | --- | --- |
| ITGAE | -1.872 | 0.002153 | integrin, alpha E (antigen CD103, human mucosal lymphocyte antigen 1; alpha polypeptide) |
| ITGAE | -1.837 | 0.001049 | integrin, alpha E (antigen CD103, human mucosal lymphocyte antigen 1; alpha polypeptide) |
| CD177 | -1.836 | 0.001049 | CD177 molecule |
| H2-DMA | -1.491 | 0.03327 | Class II histocompatibility antigen, M alpha chain |
| CCL5 | -1.442 | 0.003549 | chemokine (C-C motif) ligand 5 |
| NUP210 | -1.196 | 0.004842 | nucleoporin 210kDa |
| ITGB7 | -1.108 | 0.003194 | integrin, beta 7 |
| COTL1 | -0.94 | 0.009026 | coactosin-like 1 (Dictyostelium) |
| GVIN1 / GVINP1 | -0.8702 | 0.04398 | GTPase, very large interferon inducible pseudogene 1 |
| CD3E | -0.8288 | 0.01975 | CD3e molecule, epsilon (CD3-TCR complex) |
| MSC | -0.7953 | 0.009026 | musculin |
| ARHGAP30 | -0.7507 | 0.03716 | Rho GTPase activating protein 30 |
| GIMAP7 | -0.75 | 0.009026 | GTPase, IMAP family member 7 |
| PRMT5 | -0.6878 | 0.01187 | protein arginine methyltransferase 5 |
| NSMCE1 | -0.6086 | 0.04398 | non-SMC element 1 homolog (S. cerevisiae) |
| CX3CR1 | -0.6028 | 0.0499 | chemokine (C-X3-C motif) receptor 1 |
| DGKG | -0.5958 | 0.04583 | diacylglycerol kinase, gamma 90kDa |
| ITGB6 | 0.7005 | 0.03007 | integrin, beta 6 |
| FBXO32 | 0.7065 | 0.04807 | F-box protein 32 |
| PTPRD | 0.9848 | 0.0469 | protein tyrosine phosphatase, receptor type, D |
| STK17B | 1.037 | 0.04398 | serine/threonine kinase 17b |
| DDR1 | 1.056 | 0.03716 | discoidin domain receptor tyrosine kinase 1 |
| MS4A10 | 1.313 | 0.004361 | membrane-spanning 4-domains, subfamily A, member 10 |
| PIGT | 1.597 | 0.0499 | phosphatidylinositol glycan anchor biosynthesis, class T |
| TRAPPC3 | 1.831 | 0.009026 | trafficking protein particle complex 3 |
| PPP3R1 | 2.104 | 0.0001553 | protein phosphatase 3, regulatory subunit B, alpha |
| 2010109I03RIK | 11.22 | 9.506e-10 | Putative uncharacterized protein |

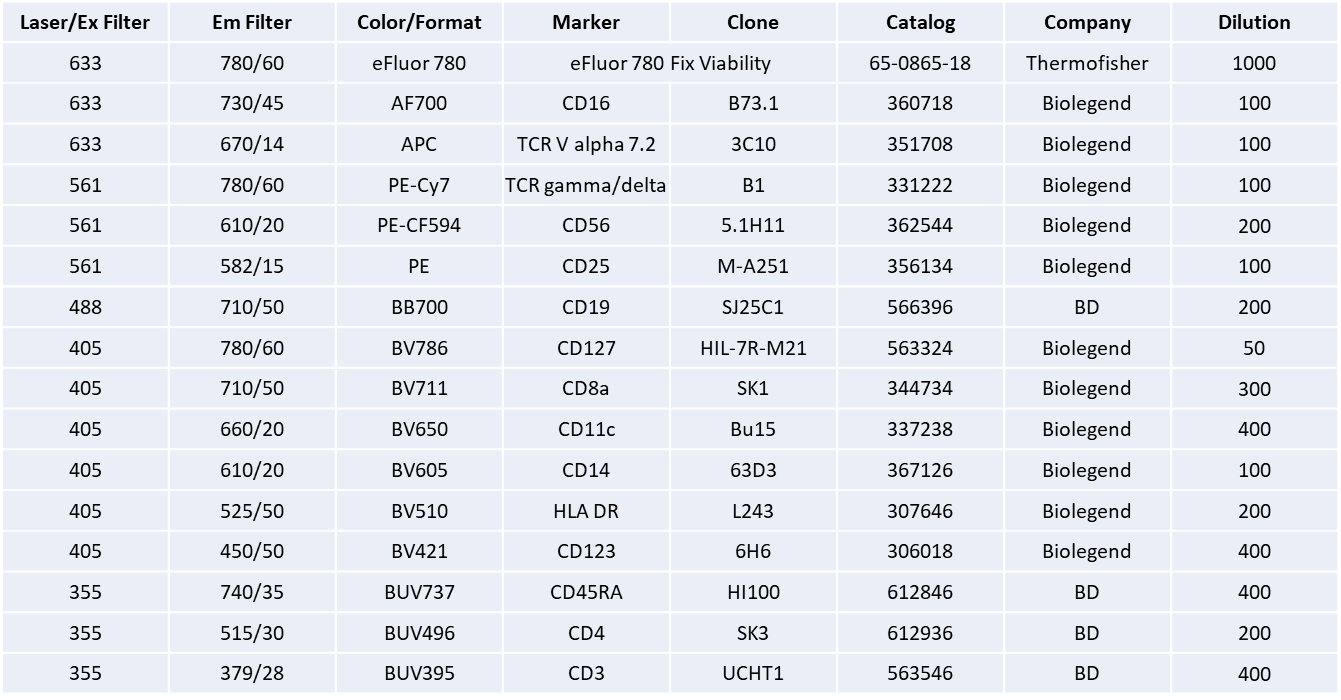
Supplementary Table 3. **Monoclonal antibody panel used to stain PBMC from UK cohort.**
